# Host specificity shapes fish viromes across lakes on an isolated remote island

**DOI:** 10.1101/2023.07.03.547585

**Authors:** Rebecca M. Grimwood, Grace Fortune-Kelly, Edward C. Holmes, Travis Ingram, Jemma L. Geoghegan

## Abstract

Fish viromes often provide insights into the origin and evolution of viruses affecting tetrapods, including those associated with significant human diseases. However, despite fish being the most diverse vertebrate group, their viruses are still understudied. We investigated the viromes of fish on Chatham Island (Rēkohu), a geographically isolated island housing 9% of New Zealand’s threatened endemic fish species. Using metatranscriptomics, we analyzed samples from seven host species across 16 waterbodies. We identified 19 fish viruses, including 16 novel species, expanding families such as the *Coronaviridae, Hantaviridae, Poxviridae*, and the recently proposed *Tosoviridae* family. Surprisingly, virome composition was not influenced by ecological factors measured, and smelt (*Retropinna retropinna*) viromes were consistent across lakes despite differences in host life history, seawater influence, and community richness. Overall, fish viromes across Rēkohu were highly diverse and revealed a long history of codivergence between host and virus despite their unique and geographically isolated ecosystem.

## Introduction

A deeper understanding of Earth’s virosphere requires a broader investigation into viruses sampled from hosts across the tree of life. Fish are a major evolutionary group that fall basal to the tetrapods. Exploring fish viromes has revealed previously hidden details about the origin and evolution of viruses found in “higher” vertebrates, including those associated with significant human disease (Chen et al., 2012; Dill et al., 2016; Grimwood et al., 2021; Lauber et al., 2017). Despite being the most diverse vertebrate group, comprising over 27,000 species (Nelson, 2006), fish are still underrepresented in virome investigations (Cobbin et al., 2021; Wille et al., 2021). In particular, we know little about how ecological factors such as community diversity, water chemistry, and life history strategy shape the viromes of fish and other aquatic vertebrates. Although some studies have begun to investigate how host ecology underpins differences in virome composition and diversity among host groups (Costa et al., 2021; Geoghegan et al., 2021; Y. Kim et al., 2017; Perry et al., 2022), studies of wild fish populations are generally limited in number and scope.

Aotearoa New Zealand is an island country located in the southwestern Pacific Ocean. Its endemic species derived from ancestral biota have been evolutionarily shaped by long periods of geographic isolation, climatic and environmental change, and dispersal after splitting from the Gondwanan supercontinent around 85 million years ago (mya) (Cooper & Millener, 1993). The Chatham Islands are an archipelago located around 800 km east of the South Island of Aotearoa New Zealand. The islands were connected to the New Zealand mainland approximately 70-65 mya, with volcanic activity responsible for the uplift of the islands from the Chatham rise less than three mya – the only part of the plateau now above sea-level (Stilwell & Consoli, 2012). The largest island is Rēkohu Wharekauri Chatham Island, hereafter Rēkohu. Rēkohu is covered by expansive wetlands, lakes and lagoons (Champion & Clayton, 2004), and hosts a rich biodiversity of aquatic and terrestrial species with high endemicity. The freshwater fish fauna includes around 9% of New Zealand’s threatened species (Roberts, 1991) as well as one endemic species. New Zealand and its surrounding islands therefore present unique, understudied ecosystems and hosts to explore questions surrounding viral diversity and evolution.

The most widespread fish species on Rēkohu is the common smelt (*Retropinna retropinna*), endemic to the islands of Aotearoa New Zealand (Ward et al., 2005; Woods, 1968). Smelt are normally diadromous, spawning in coastal estuaries and lowland rivers with larvae migrating to rear in the ocean and returning as postlarval ‘whitebait’. However, smelt readily landlock when they reach isolated lakes, completing their lifecycle within the lake. Rēkohu is home to diadromous smelt in coastal streams and the large lagoon Te Whanga, as well as numerous lacustrine (landlocked) populations. Smelt also occur in both freshwater and salt-influenced lakes, with a range of coexisting fish species richness, offering an opportunity to investigate impacts of life history and environmental variation on virome composition.

Here, we sampled seven fish species, including smelt, from 16 waterbodies across Rēkohu, and used metatranscriptomics to reveal the diversity and distribution of their viruses. Factors such as life history (diadromous or lacustrine), fish community richness, and seawater influence, were explored to determine how these factors may shape fish viromes. This study expands on an initial screen of smelt from Rēkohu (Perry et al., 2022) and highlights novel viruses of additional species coexisting in these environments.

## Methods

### Animal ethics

Animal capture and euthanasia were approved by the University of Otago Animal Ethics Committee (AUP-21-93).

### Fish tissue collection and storage

Gill tissue was sampled from seven fish species across 16 waterbodies on Rēkohu (43.9271° S, 176.4592° W) in November 2021 (Table 1 and Figure 1). As the gills are a point of entry and site of replication for many fish-associated viruses (Kim & Leong, 1999), they were chosen as the tissue type of interest for this study. Smelt (*R. retropinna*) were sampled from all 16 sites. At five lake sites, all other fish species present were also sampled, including *Anguilla australis* (shortfin eel), *Anguilla dieffenbachii* (longfin eel), *Galaxias maculatus* (inanga), *Neochanna rekohua* (mudfish), *Galaxias argenteus* (giant kōkopu), and *Forsterygion nigripenne* (estuarine triplefin), each sampled from between one and three of the waterbodies (Table 1).

**Table 1.**
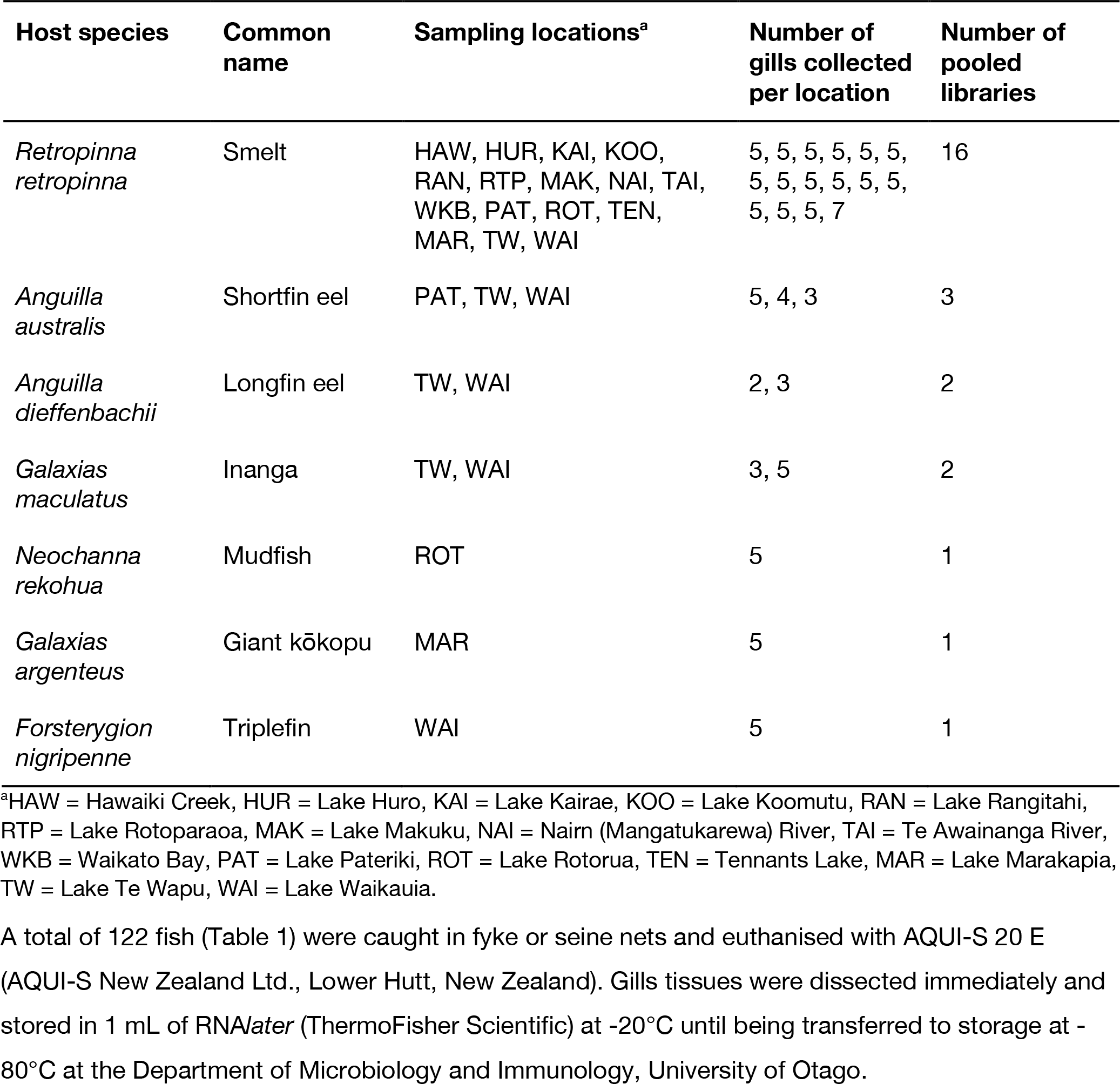
Overview of host species and sampling locations.

**Figure 1.**
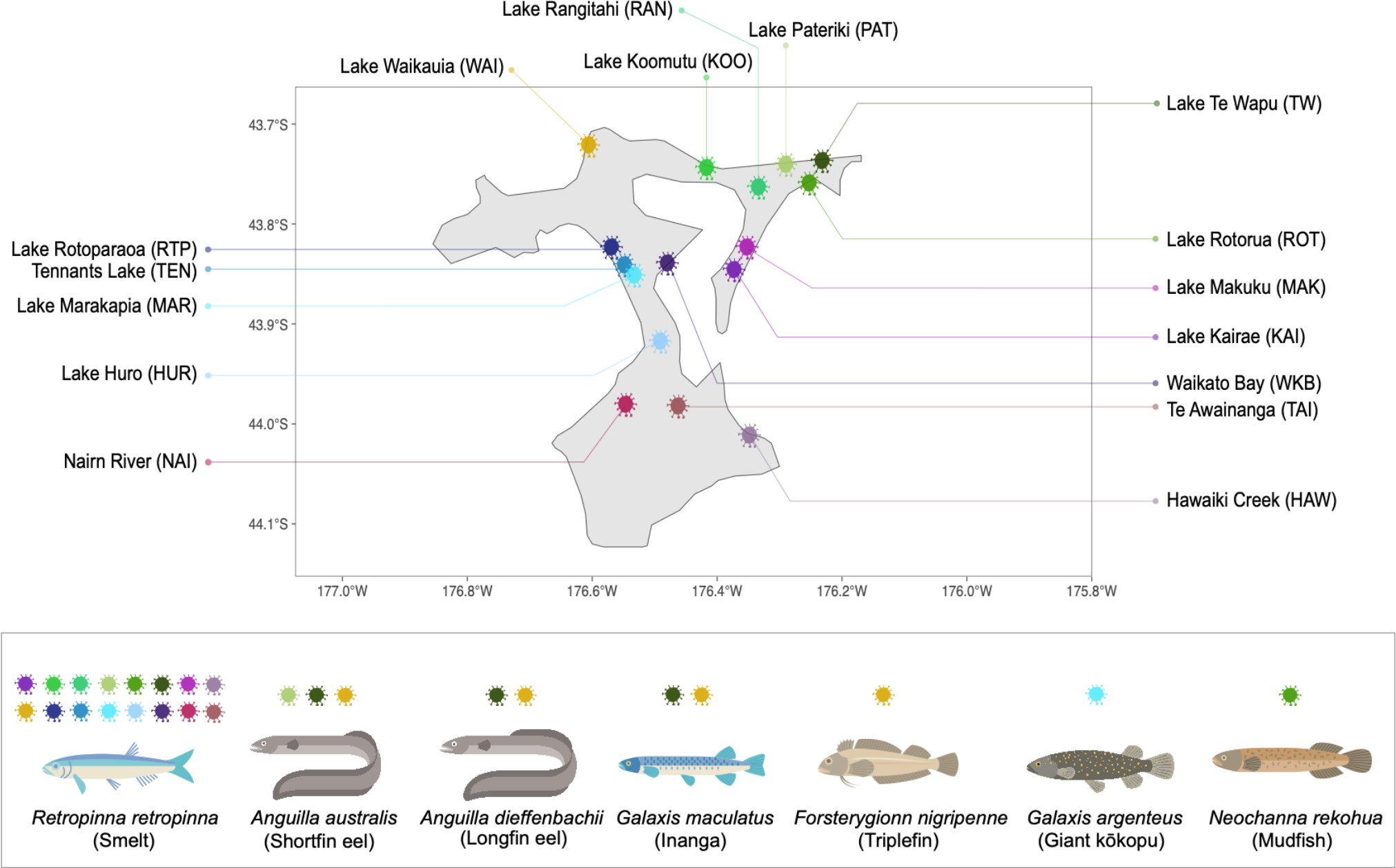
Map of Rēkohu showing the 16 sampling sites (waterbodies) of the seven fish species under investigation. Sites where each species was sampled are indicated as coloured virus symbols on the map and corresponding symbols above each species. Sampling locations ranged from one to 16 sites per species.

A total of 122 fish (Table 1) were caught in fyke or seine nets and euthanised with AQUI-S 20 E (AQUI-S New Zealand Ltd., Lower Hutt, New Zealand). Gills tissues were dissected immediately and stored in 1 mL of RNA*later* (ThermoFisher Scientific) at -20°C until being transferred to storage at - 80°C at the Department of Microbiology and Immunology, University of Otago.

### RNA extraction and sequencing

Frozen gill tissue was partially thawed and ∼10-15 mg was taken from each sample using sterilised scalpels and forceps and placed into RLT lysis buffer containing 1% ß-mercaptoethanol and 0.5% (v/v) Reagent DX. Dissected tissues were homogenised in the lysis buffer using a Qiagen TissueRupture for around 30 seconds each. The resulting homogenates were centrifuged to remove residual tissue and the RNeasy Plus Mini Kit (Qiagen) protocol was followed to extract whole RNA. An additional 500 μL ethanol wash and 1 minute spin at ≥8000 x *g* (≥10,000 rpm) was added before the final RNA elution step to prevent guanidine contamination.

RNA was quantified using a NanoDrop Spectrophotometer (ThermoFisher) and then pooled by host species and location for a total of 26 pools (Table 1) each comprising RNA from 2-7 individuals. 10-15 μL of total RNA per individual was pooled. Pooled RNA was sent for total RNA sequencing on the Illumina NovaSeq 6000 platform. Illumina Stranded Total RNA Prep with Ribo-Zero plus (Illumina) was used for library preparation and 150 bp paired-end reads were generated.

### Transcriptome assembly, annotation, and virus screening

Paired-end reads were assembled *de novo* using Trinity RNA-seq (v2.11) (Grabherr et al., 2011) with default parameters, plus the ‘trimmomatic’ flag option to perform pre-assembly quality trimming. Resulting contigs were annotated based on similarity searches at both nucleotide and amino acid levels to identify potential viral sequences. The BLASTn algorithm (Camacho et al., 2009) was used to screen contigs against NCBI’s nucleotide (nt) database and DIAMOND BLASTx (v2.02.2) (Buchfink et al., 2021) was used for screening against the non-redundant (nr) protein database. Any sequences predicted to be viral were manually checked with additional BLASTx and BLASTn searches using the online BLAST server (https://blast.ncbi.nlm.nih.gov/Blast.cgi) to eliminate false positive hits. Putative viruses with e-values of greater than 1×10^-10^ or where the top BLAST hits included non-viral species were excluded from further analysis. Segments of DNA viruses were also subjected to additional nucleotide BLAST searches to check for possible homology to fish genomic DNA and exclude endogenous viral elements.

### Virus phylogenetic analysis

Evolutionary relationships of the viruses identified in the gill meta-transcriptomes were explored using conserved protein segments, usually the RNA-dependent RNA polymerase (RdRp), DNA polymerase, or LO7 (a hexon-like protein) in the case of viruses from *Adomaviridae* (Iwanowicz et al., 2020). Viral amino acid sequences identified as likely being vertebrate-associated based on hosts species assignments of their closest relatives and that could be assigned to at least order or family level were aligned with their top BLAST hit and a representative range of viruses from different host species in their respective taxonomic orders collected from NCBI Taxonomy (https://www.ncbi.nlm.nih.gov/taxonomy) using MAFFT (v7.450) (Katoh & Standley, 2013). The L-INS-i algorithm was used for all alignments, with the exception of the *Flaviviridae* for which the E-INS-i algorithm was used to ensure alignment of the conserved glycine-aspartic acid (GDD) RdRp active site motif (O’Reilly & Kao, 1998). Alignments were visualised in Geneious Prime (v2020.2.4) (https://www.geneious.com) and ambiguously aligned regions were trimmed using trimAL (v1.2) (Capella-Gutiérrez et al., 2009) with the ‘automated1’ flag. IQ-TREE (v.1.6.12) (Nguyen et al., 2015) was used to estimate maximum likelihood trees for each family under consideration. The LG amino acid substitution model was used with 1000 ultra-fast bootstrapping replicates (Hoang et al., 2018) and the ‘alrt’ flag enabled to perform 1000 bootstrap replicates for the SH-like approximate likelihood ratio test (Guindon et al., 2010). Phylogenetic trees were annotated in FigTree (v1.4.4) (http://tree.bio.ed.ac.uk/software/figtree/) and rooted at their midpoints.

### Polymerase alignments

Amino acid alignments of the virus polymerase sequences were created to complement phylogenetic trees and illustrate which segments for each virus were identified in the sequencing read data. Closely related viral sequences for each virus found in this study were collected as references and aligned in Geneious Prime (v2020.2.4) using the L-INS-I algorithm within MAFFT (v7.450). If the same species was found across multiple locations with different top BLAST hits, one of the top hits (usually the longest or most complete) was chosen and all instances of the virus were aligned to it. Unaligned ends were manually trimmed for pictorial clarity and relevant catalytic sites or motifs were highlighted if this information was available for the reference sequences.

### Virome abundances

To estimate the abundance virus transcripts, assembled contigs were quantified using the ‘align and estimate’ module within Trinity RNA-seq with the ‘prep reference’ flag set. RNA-seq by Expectation-Maximization (RSEM) (Li & Dewey, 2011) was used as the abundance estimation method and Bowtie 2 (Langmead & Salzberg, 2012) as the alignment method. Transcript abundances were standardised by dividing by respective sequencing library depths. To reduce incorrect assignment of viruses to libraries due to index hopping, viruses with a read count less than 0.1% of the highest count for that virus across the other libraries were considered contamination and omitted from that library. Virome abundances (both total and vertebrate-associated) were compared alongside the standardised abundance of the stably expressed host gene *40S Ribosomal Protein S13* (RPS13). RPS13 shows little inter- and intra-tissue expression level variation making it a suitable reference gene for transcript expression comparisons (Robichaux et al., 2016). For heatmaps, standardised abundances were further normalised across viral family before plotting by calculating the proportion of a virus family in a sample out of the total abundance of that family across all libraries.

### Ecological analysis

The diversity of all fish species and smelt-only viromes were explored using R (v4.1.1). Alpha diversity was measured as observed richness and Shannon diversity indices of normalised abundances. Welch two sample t-tests were performed to compare alpha diversity measures between diadromous and landlocked smelt populations.

The relationship between vertebrate-associated virus families and various ecological factors was explored for all fish species, and for smelt-only pools. These factors were seawater influence, life history, and fish community richness. Seawater influence at each site was the only factor investigated including all fish species in the study. Each site was characterised based on the seawater influence (fresh, brackish, or seawater) expected at the rearing environment for larval smelt: in the lake for landlocked populations or in the ocean or Te Whanga for diadromous populations. Life histories were either landlocked or diadromous smelt populations, with landlocking assessed based on the present connectivity of the lake and/or genetic evidence of isolation from diadromous populations (Ara et al., unpub. data). Community complexity (richness) in each lake was coded as 0, 1-2, 3-4, or >4 fish species (in addition to smelt) at a site, based on previous sampling.

Vertebrate family-level virome abundances were normalised and a distance matrix was created using the vdist function available in the vegan package (Oksanen et al., 2022) with Bray-Curtis dissimilarity as the distance measure. The distance matrix was then used to perform multivariate ordination using non-metric multidimensional scaling (NMDS) on the distance matrix with the metaMDS function from vegan. NMDS data points were plotted and coloured by the corresponding ecological factor being investigated using ggplot2 (Wickham, 2016). Permutational multivariate analysis of variance (PERMANOVA) using the vegan function adonis2 was used to test for statistical significance of the effect of each ecological factor on virome composition. In the case of seawater influence on the viromes of the full host set, the significance of seawater influence both with and without the consideration of host species was tested.

Virome heterogenity among sites, measured as the multivariate dispersion of smelt virome compositions, was also compared to the three ecological factors tested (seawater influence, community richness, and life history). Dispersion was compared among groups using the distance matrix and the betadisper function in vegan, and statistical significance was tested using the anova function.

Full R code detailing all packages and versions used, and formatted data used to produce associated results are available on GitHub (see Data availability and reproducibility).

### Virus nomenclature

Viruses identified were considered novel if they shared less than 90% sequence identity with their most closely related virus (ICTV, 2012). Novel viruses were named according to their host species, either using the genus name or a common name where more distinction between several related hosts was needed.

### Data availability and reproducibility

Raw sequencing data are available under BioProject *pending* while viral sequences identified are under GenBank accession numbers *pending*. Please see GitHub for the full R code, packages, and versions used for analyses and accompanying abundance data and meta/site data: https://github.com/maybec49/Rekohu_viromes.

## Results

### Metatranscriptomic sequencing

Gill tissue RNA from seven fish hosts was pooled into 26 libraries covering both species and sampling locations to identify any viruses present in these hosts. Metatranscriptomic sequencing yielded 46.4 – 86.6 million 150 bp paired-end reads per library. A combined total of 2,857,823,201 reads were obtained. No quality issues were identified in any of the libraries. Between 393,574 and 827,518 contigs were assembled per library using Trinity RNA-seq. Viral abundances ranged from 0.000047% and 0.024% of total standardised abundances for each library.

### Overview of viromes and abundances

At least one likely vertebrate host-associated virus was identified in 19 of the 26 libraries (range: 0 to 5 viruses). No vertebrate host-associated viruses were found in triplefin, mudfish, or giant kōkopu. Overall, 19 virus species (38 individual viruses) were identified across 13 viral taxonomic families or orders (Figure 2). Of the virus species found, 16 were novel (sharing less than 90% amino acid identity with their most closely related virus species). Vertebrate host-associated viruses made up 0 to 99.2% of the total virome abundance in each library. The virome abundances of all the libraries were lower than that of host RPS13 expression, except for the total virome abundance in longfin eels from Lake Te Wapu (Figure 2).

**Figure 2.**
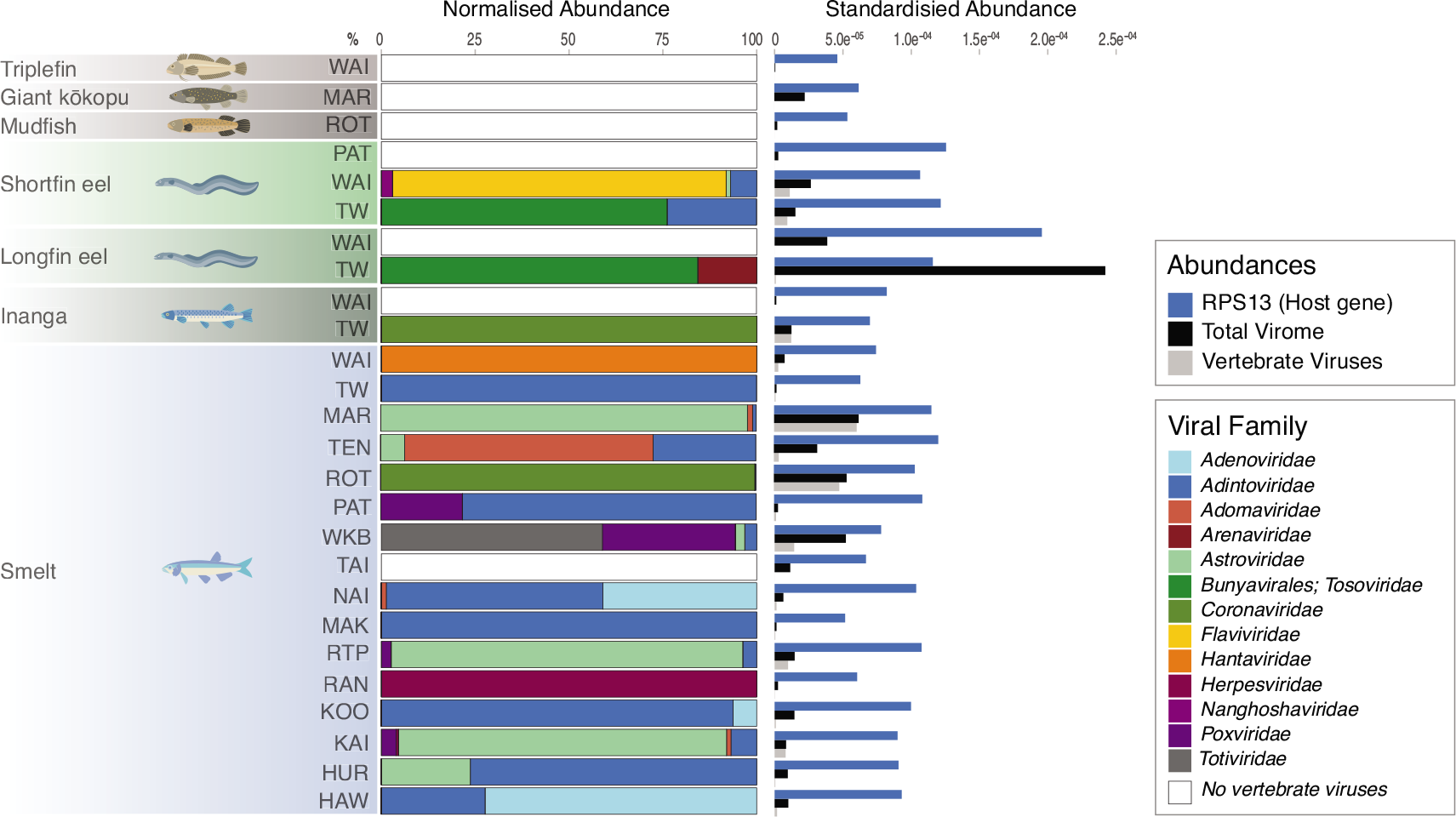
Compostion of virome abundances for each sequencing library. Normalised abundance (%) of the vertebrate-associated virus families in each of the 26 libraries (left) and standardised abundances of the RPS13 host gene expression (blue), total viromes (black), and vertebrate-associated viruses (grey) (right).

Viruses from families such as the *Adintoviridae*, *Adomaviridae*, *Astroviridae* and *Poxviridae* were the most widely distributed and abundant viral families among smelt (Figure 3). In comparison, viruses from families such as the *Arenaviridae*, *Hantaviridae*, *Flaviviridae* and *Nanghoshaviridae* were only found in single samples, primarily shortfin and longfin eels. Overall, smelt viromes were mostly homogenous with a small number of viruses from additional, less abundant families present in a few smelt samples. Samples from freshwater lakes had an average of 1.8 vertebrate-associated viral families, while those from seawater had an average of 1.7 families. Sampling locations with a brackish influence contained 1.1 families on average.

**Figure 3.**
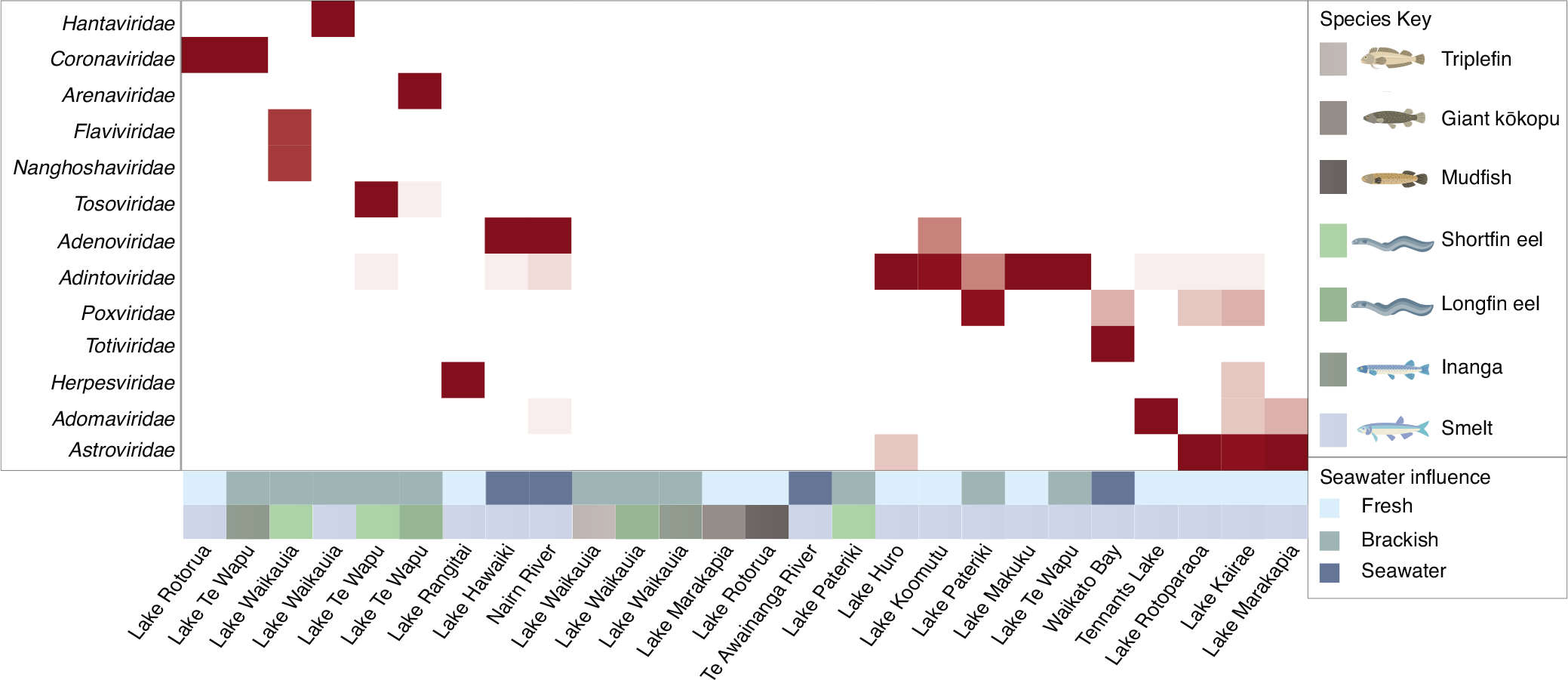
Heatmap of standardised abundances of viral families per library. Viral families (rows) and libraries (columns) were both clustered according to Bray-Curtis distances. Abundances were normalised by viral family. Darker red blocks indicate higher abundance. Seawater influence in each lake (top) and host species (bottom) shown as coloured blocks below heatmap.

### DNA viruses discovered in Rēkohu fish

Viruses from five double-stranded DNA viral families were identified in smelt hosts and seven viral species in four of these families (*Adenoviridae*, *Adintoviridae*, *Adomaviridae*, and *Poxviridae*) could be subjected to phylogenetical analysis. Various partial segments from viruses in the *Herpesviridae* were identified, including partial capsid proteins. However, due to the lack of sufficient DNA polymerase sequences these were assessed phylogenetically. All DNA viruses assessed clustered in distinct fish virus-associated phylogenetic clades.

### Adintoviridae

Type B DNA polymerase (PolB) segments from an *Adintovirus*, *Porure adintovirus-1*, previously associated with smelt on Rēkohu (Perry et al., 2022) were identified in 12 of the 16 waterbodies (Figure 4). *Porure adintovirus-1* in each of these samples shared a minimum of 97% nucleotide and amino acid identity with the original sequence of this virus found in 2022 (accession: UNJ19240.1), such that the viruses found in these additional sampling sites are all the same species. Adintoviruses represent a recently proposed group of exogenous and endogenous polinton-like dsDNA viruses (Starrett et al., 2021). While many adintoviruses have been endogenised into host genomes and subsequently degraded by frameshift and nonsense mutations, the recovery of segments covering a DNA polymerase without evidence of such degradation suggests these sequences to be from an exogenous adintovirus species. Furthermore, the original PolB sequence had a length of 155 amino acids, while the sequences found in this study ranged from 134 to 276 amino acids, expanding the known sequence spanning the PolB segment of this virus (Figure 4). All fish-associated viral species in this family formed a distinct clade in the phylogeny.

**Figure 4.**
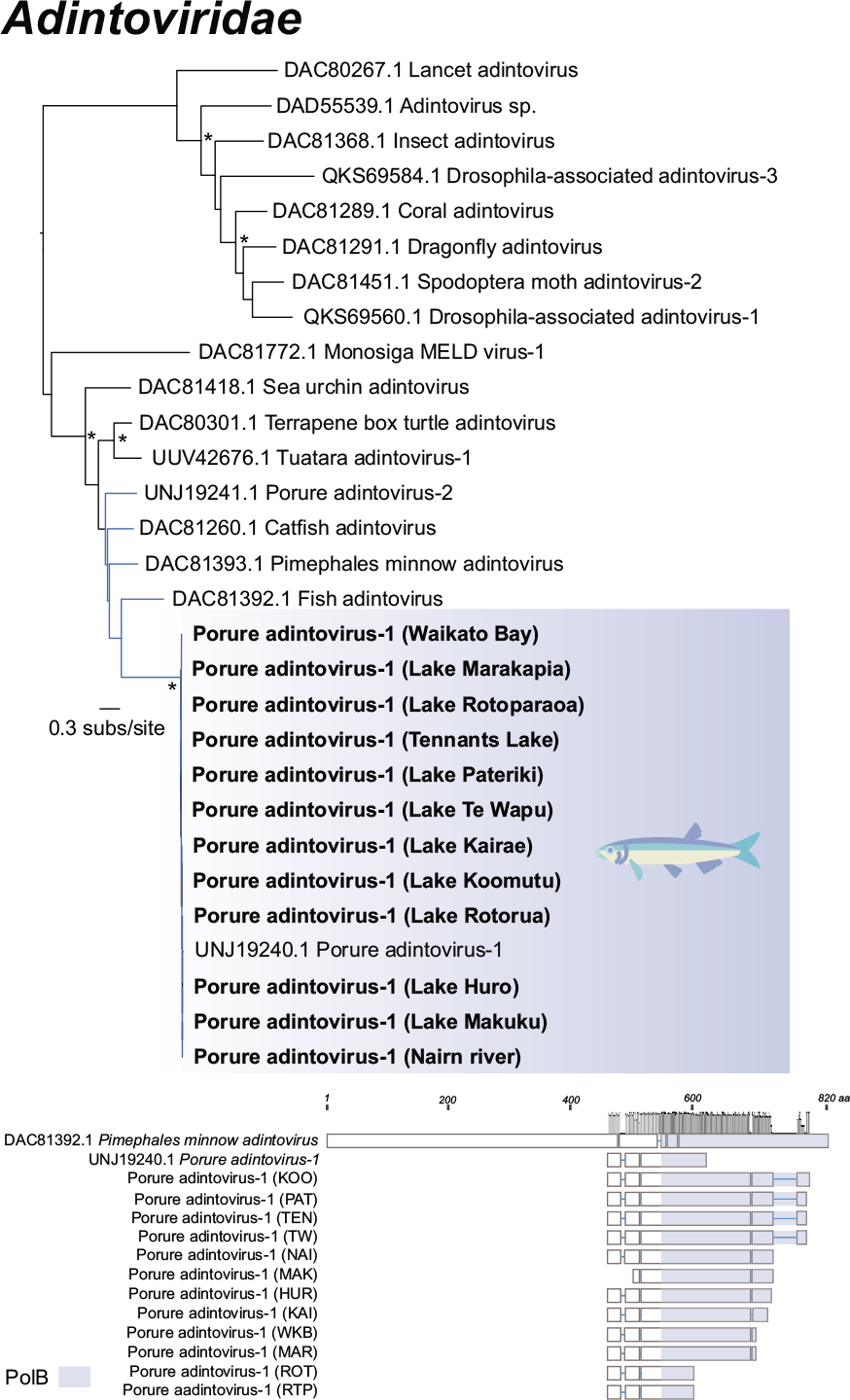
Maximum likelihood tree of PolB segments of Porure adintovirus-1 within the Adintoviridae (dsDNA) and recovered segments (bottom; amino acids). Smelt viruses identified in this study are bolded and highlighted in blue. Fish-associated viruses are indicated by blue branches. UF-bootstrap node support values ≥ 95% are denoted by an asterisk (*) and the tree is rooted at its midpoint. The height of grey bars above conserved regions of the PolB segments indicates percentage identity.

### Adenoviridae

Two novel adenoviruses, Retropinna adenovirus-1 and Retropinna adenovirus-2, were discovered in smelt from Hawaiki Creek, Lake Koomutu, and Nairn River (Figure 5a). Retropinna adenovirus-1 shared 47.2-47.9% amino acid identity with *Raptor adenovirus* (accession: UTK57358.1; e-value: 2×10^-16^) and *Raptor adenovirus-1* (accession: YP_004414800.1; e-value: 8×10^-27^), while Retropinna adenovirus-2 shared ∼47.6% amino acid identity to *Gouldian finch adenovirus-1* (accession: AGU01648.1; e-value: 2×10^-45^). Capsid protein segments from both viruses were also identified (see Supplementary Table S2). Although some adenoviruses can cause disease in avian hosts, the closest relatives of the novel adenoviruses have not been definitively linked to disease in their hosts (Kovács & Benkő, 2009; Rinder et al., 2020). Phylogenetically, the two viruses cluster with *White sturgeon adenovirus-1*, forming a distinct clade of adenoviruses associated with cold-blooded vertebrate hosts (Figure 5a). Notably, *White sturgeon adenovirus-1* has been found to cause lesions (C. H. Kim & Leong, 1999).

**Figure 5.**
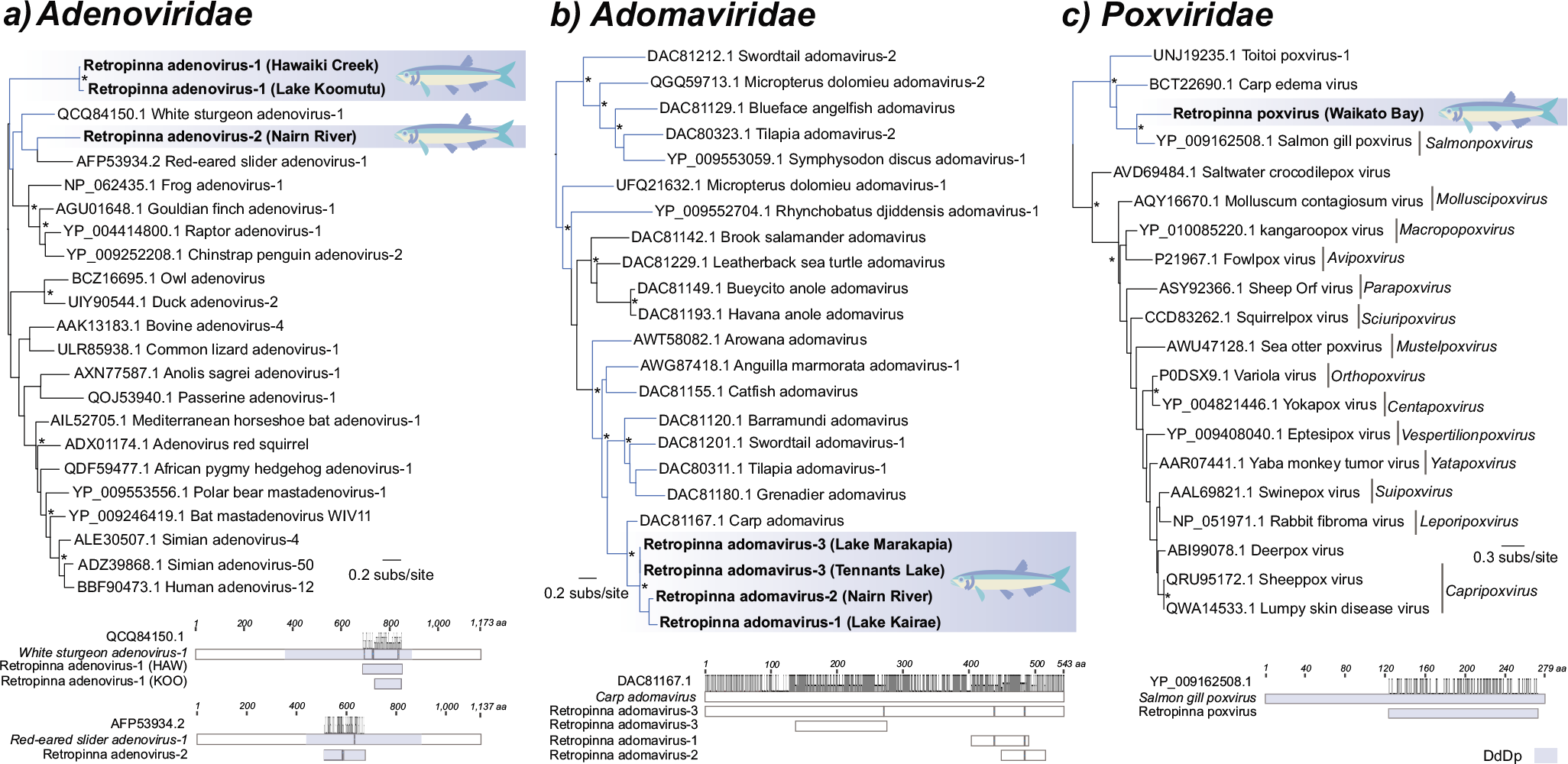
Maximum likelihood trees of novel dsDNA viruses identified in this study (top) and respective illustrations of recovered segments (amino acids) shown under that of closely related viruses (bottom). Fish-associated viruses are denoted by blue branches. Smelt viruses identified in this study bolded and highlighted in blue. UF-bootstrap node support values ≥ 95% are indicated by an asterisk (*). All trees were rooted at their midpoint. The height of the grey bars above conserved regions indicate percentage identities. A) Phylogenetic tree and polymerase structures of the DdDp segments of novel viruses identified within the *Adenoviridae*. B) Phylogenetic tree and sequence organisations of the LO7 segments of novel viruses identified within the *Adomaviridae*. C) Phylogenetic tree and polymerase organisation of the DdDp segment of a novel poxvirus within the *Poxviridae*.

### Adomaviridae

Segments of the LO7 (hexon-like) protein from three novel adomaviruses in smelt were found, sharing 66.7-75.8% acid identity with *Carp adomavirus* (accession: DAC81167.1; e-values: 2×10-^20^-0). Retropinna adomavirus-1 was found in Lake Kairae, Retropinna adomavirus-2 in Nairn River, and Retropinna adomavirus-3 in Lake Marakapia and Tennants Lake. These all clustered within a large group of other fish-associated viruses (Figure 5b). Additional sequences from early proteins EO2-4 and the late proteins LO5, 6 and 8 were recovered from Retropinna adomavirus-3 (Supplementary Table S2). Although the *Adomaviridae* is a relatively emergent family of non-enveloped viruses with similarities to those in the *Adenoviridae* (Welch et al., 2020), at least one virus within this family has been associated with skin disease in fish (Dill et al., 2018).

### Poxviridae

Multiple segments, including a DNA polymerase and capsid proteins, from a novel poxvirus were identified in smelt from Waikato Bay (see Supplementary Table S2). A 150 amino acid segment of the 18 kDa subunit of an DNA-directed RNA polymerase (RPO18) was used for phylogenetic analysis (Figure 5c). The virus, named Retropinna poxvirus, shared 56.4% amino acid identity with the RPO18 from *Salmon gill poxvirus* (genus: *Salmonpoxvirus*; accession: YP_009162508.1; e-value: 3 x10^-46^). The virus falls within a basal fish virus-associated clade. Poxvirus-associated diseases are less well-studied in fish due to the inability of the viruses to be propagated in cell culture, but diseases such as Salmon gill poxvirus, Carp edema and koi sleepy disease can cause mortalities up to 70-80% (Gjessing et al., 2016).

### RNA viruses in Rēkohu fish

A further 12 virus species from eight RNA families were identified in fish from Rēkohu lakes. This included viruses with double-stranded (ds), positive (+)-, negative (-)-, and ambi-sense (+/-) single-stranded (ss) RNA genomes. Viruses were found across smelt, shortfin and longfin eel, and inanga fish hosts.

### Arenaviridae, Hantaviridae and Totiviridae

Nucleoprotein, glycoprotein, and L segments from a novel arenavirus (+/- ssRNA) were identified in Lake Te Wapu longfin eels (see Supplementary Table S2 and Figure 6b). The virus, named Longfin eel arenavirus, shared 48.8% amino acid identity to *Salmon pescarenavirus-2* (accession: QEG08232.1; e-value: 3×10^-19^). Of note is that the segment recovered did not overlap the conserved RdRp region making its position in the *Arenaviridae* less certain (Figure 6a). Arenaviruses were originally thought to be confined to mammals, but have since been found in cold-blooded vertebrates, including in association with disease pathology and symptoms in salmon (Mordecai et al., 2019a).

**Figure 6.**
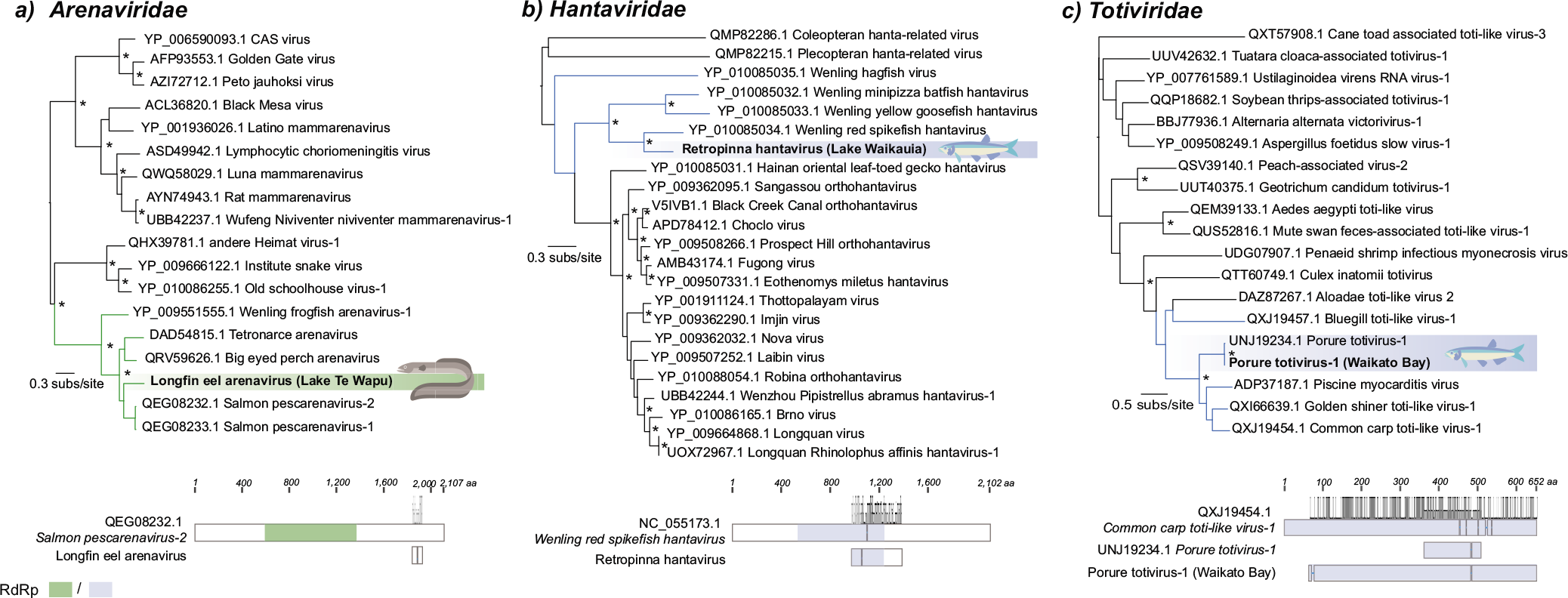
Maximum likelihood trees of viruses in Arenaviridae (+/- ssRNA), Hantaviridae (-ssRNA), and Totiviridae (dsRNA) identified in this study (top) and recovered RdRp segments (amino acids) shown against that of a representative virus (bottom). Viruses identified in this study bolded and highlighted in blue for smelt- and green for eel-associated viruses. Fish-associated viruses indicated by blue or green branches. UF-bootstrap node support values ≥ 95% are denoted by an asterisk (*). All trees were rooted at their midpoint. The height of grey bars above conserved regions indicates percentage identities. A) Phylogenetic tree and structure of the RdRp segment of a novel longfin eel virus within the *Arenaviridae*. B) Phylogenetic tree and structure of the RdRp segment of a smelt virus identified within the *Hantaviridae*. C) Phylogenetic tree and structure the RdRp of Porure totivtus-1 from Waikato Bay within the *Totiviridae*.

In addition, glycoprotein, nucleoprotein, and RdRp segments of a novel hantavirus (-ssRNA) was found in smelt from Lake Waikauia (Figure 6b and Supplementary Table S2). This new virus was most closely related to *Wenling red spikefish hantavirus* (identity percentage: 50.3%; accession: YP_010085034.1; e-value: 9×10^-142^). The *Hantaviridae* were also previously dominated by mammalian-associated viruses, with more recently identified viruses in fish (Shi et al., 2018).

*Porure totivirus-1* (dsRNA) has been previously identified in smelt (Perry *et al*. in 2022) and was identified here in smelt in Waikato Bay (Figure 6c). A 579-amino acid segment of the RdRp was recovered, more than doubling the known sequence length of the original 144 amino acid residue sequence described previously (Perry *et al*. in 2022; accession: UNJ19234.1). While totiviruses were thought to primarily infect unicellular organisms, such as fungi, there is growing evidence for infection of invertebrates, mammalian cell lines, and even association with cardiomyopathy syndrome in farmed Atlantic salmon (Løvoll et al., 2010).

All viruses identified in Rēkohu from the *Arenaviridae*, *Hantaviridae* and *Totiviridae* fell confidently into fish-associated virus clades.

### Astroviridae, Coronaviridae and Flaviviridae

Several highly divergent (+)ssRNA viruses were found from three families: *Astroviridae, Coronaviridae* and *Flaviviridae*. All the novel viruses in these families, with the exception of Retropinna-associated astrovirus, grouped with other fish or cold-blooded vertebrate viruses (Figure 7).

**Figure 7.**
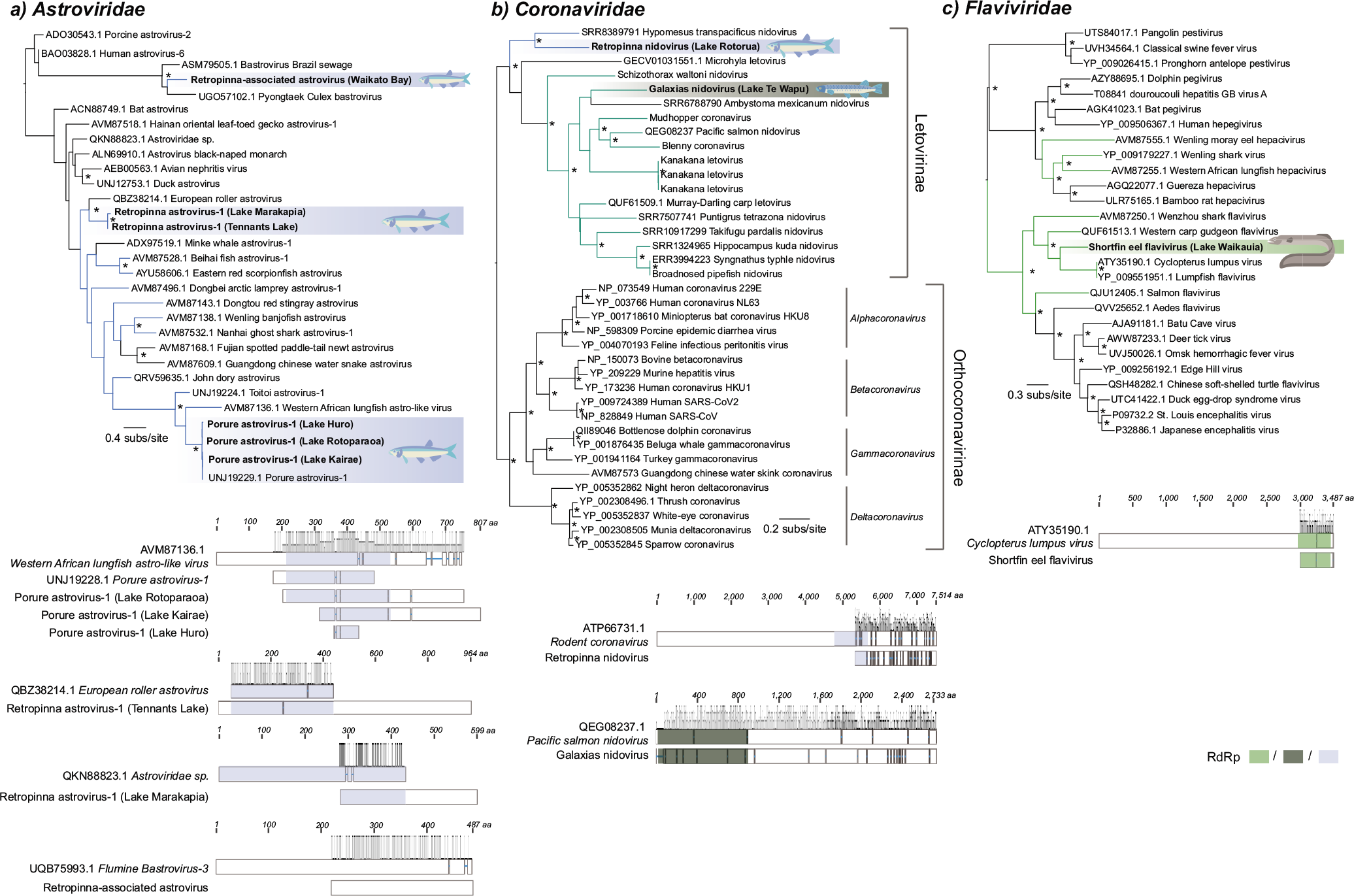
Maximum likelihood tree of (+) ssRNA viruses identified in this study (top) and recovered segments (amino acids) shown against that of closely related virus (bottom). Viruses identified in this study are bolded and highlighted in blue for smelt-, green for eel-, and dark green for inanga-associated viruses. Fish-associated viruses indicated by blue or green branches. UF-bootstrap node support values ≥ 95% denoted by an asterisk (*). All trees were rooted at their midpoint. The height of grey bars above conserved regions indicate percentage identities. A) Phylogenetic tree and organisations of the RdRp fragments of viruses identified within the *Astroviridae*. B) Phylogenetic tree and organisations of the RdRp segments of novel viruses identified within the *Coronaviridae*. C) Phylogenetic tree and organization of a polyprotein of a novel flavivirus within the *Flaviviridae*.

Both novel and previously described astroviruses were identified. *Porure astrovirus-1* was found in smelt from lakes Huro, Rotoparaoa, and Kairae, sharing 98.5-100% amino acid identity with the original virus (accession: UNJ19228.1). Like the other previously known Rēkohu smelt viruses found here, the known sequence of this original virus was also expanded (towards the 3’ end) by sequences recovered here (Figure 7a). Full length transcripts from a novel virus, Retropinna astrovirus-1, were found in smelt from Tennants Lake and Lake Marakapia (see Supplementary Table S2 and Figure 7a). The virus was most closely related to *European roller astrovirus* (percentage identity: 50.0%; accession: QBZ38214.1; e-value: 2×10^-123^) and an *Astroviridae sp.* (percentage identity: 41.7%; accession: QKN88823.1; e-value: 2×10^-24^), respectively. Finally, another *astrovirus* from Waikato Bay, Retropinna-associated astrovirus was identified and most closely related to *Flumine bastrovirus-3* (percentage identity: 54.7%; accession: UQB75993.1; e-value: 1×10^-72^). This virus clustered with invertebrate host-associated and environmental viruses, suggesting it is unlikely to be directly infecting smelt but rather a dietary or environmental contaminant (Figure 7a).

Two novel viruses falling in the *Letovirinae* subfamily of the *Coronaviridae* were identified in inanga and smelt (see Supplementary Table S2 and Figure 7b). Galaxias nidovirus, found in Lake Te Wapu from inanga, was related to *Pacific salmon nidovirus* (percentage identity: 48.4%; accession: QEG08237.1; e-value: 0.0), while the Retropinna nidovirus from smelt in Lake Rotorua was most similar in sequence to *Rodent coronavirus* (percentage identity: 37.5%; accession: ATP66731.1; e-value: 0.0) but clustered phylogenetically with another divergent virus that was previously identified in a gill transcriptomes of delta smelt (*Hypomesus transpacificus*) in the San Francisco Estuary, California, USA (SRA accession: SRR8389791) (Komoroske et al., 2021) using the Serratus virus discovery tool (Edgar et al., 2022).

Finally, a partial putative envelope protein and a 510-amino acid segment of an RdRp from a novel flavivirus was recovered from Lake Waikauia shortfin eel (see Supplementary Table S2 and Figure 7c). This flavivirus shared 47.4% amino acid identity with *Cyclopterus lumpus virus* (accession: ATY35190.1; e-value: 3×10^-129^), a virus which has been associated with, but not confirmed as the cause of, disease with high mortality in young *C. lumpus* fish (Skoge et al., 2018).

### Highly divergent viruses identified in longfin and shortfin eels: Nanghoshaviridae and Tosoviridae

Three highly divergent viruses with initially uncertain viral family placement were identified in several shortfin and longfin eel samples (Figure 8).

**Figure 8.**
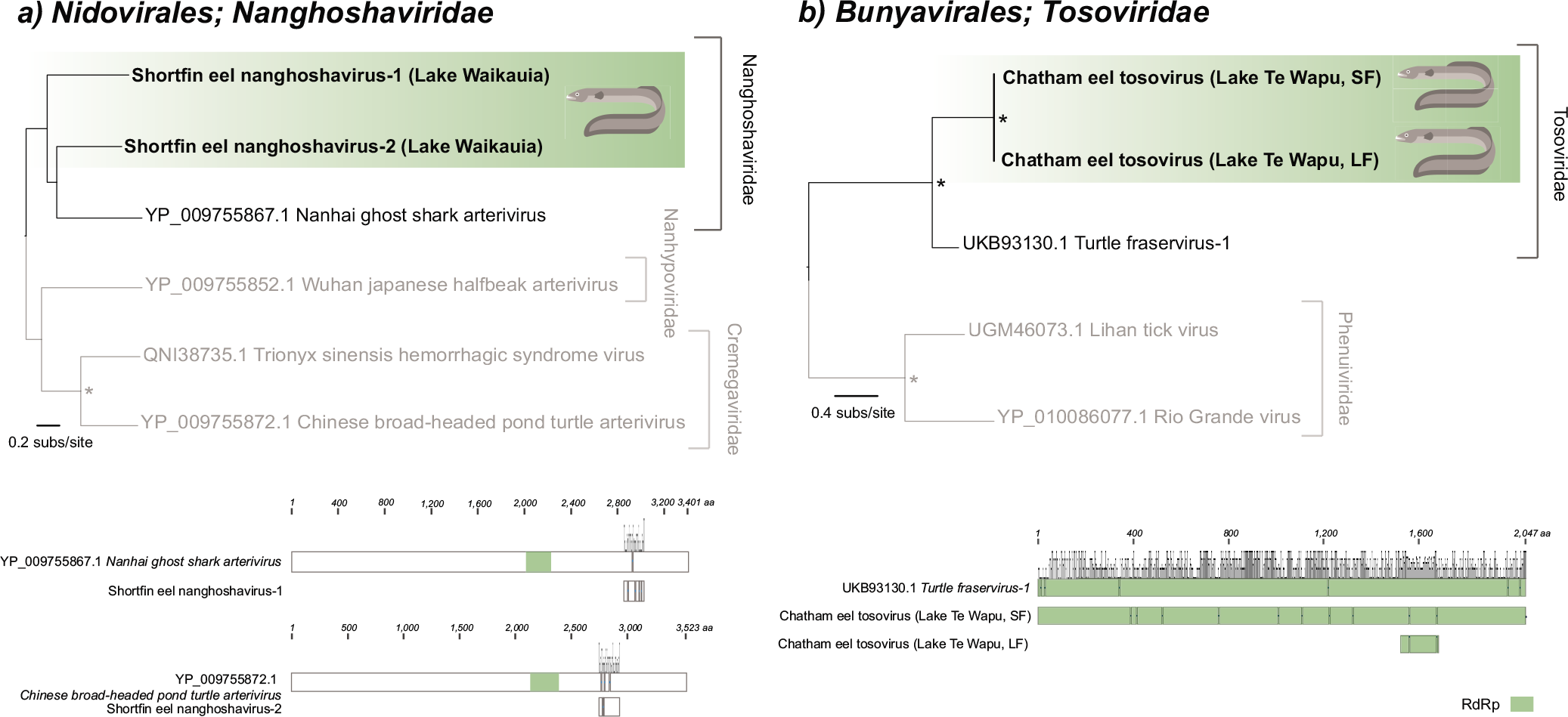
Maximum likelihood trees of novel viruses within the Nidovirales (+ ssRNA) and Bunyavirales (-ssRNA) (top) and recovered segments (amino acids) (bottom). RdRp organisations are shown alongside a closely related virus reference. The height of grey bars above conserved regions indicates percentage identities. Eel viruses identified in this study bolded and highlighted in green. Fish-associated viruses indicated by green branches. UF-bootstrap node support values ≥ 95% are denoted by an asterisk (*). All trees were rooted at their midpoints. A) Phylogenetic tree and organisation of the novel virus within the *Nanghoshaviridae*, with the *Nanhypoviridae* and *Cremegaviridae* utilized as outgroups (greyed-out branches). B) Phylogenetic tree and organisation of the novel tosovirus within the *Tosoviridae*, with the *Phenuiviridae* as an outgroup (greyed-out branches).

Two viruses from Lake Waikauia shortfin eels shared significant amino acid identity with viruses from the *Nidovirales* order (Figure 8a). The closest BLAST hit for Shortfin eel nanghoshavirus-1 was *Nanhai ghost shark arterivirus* (percentage identity: 34.9%; accession: YP_009755867.1; e-value: 9×10^-12^), while the top hit for Shortfin eel nanghoshavirus-2 was the *Chinese broad-headed pond turtle arterivirus* (percentage identity: 33.9%; accession: YP_009755872.1; e-value: 2×10^-14^). When subjected to phylogenetic analysis both viruses fell within the *Nanghoshaviridae,* a family comprising fish-associated viruses found in China (Shi et al., 2018). However, since the recovered segments do not overlap the conserved RdRp and were recovered from the same library, it is possible that these two viruses represent different regions of ORF1ab from a single virus.

Full-length nucleoprotein and RdRp transcripts, as well as a near full-length glycoprotein transcript from a highly divergent virus was discovered in both longfin and shortfin eels from Lake Te Wapu, sharing 99.8% nucleotide sequence identity and indicating it as being the same species in both hosts (Supplementary Table S2). Phylogenetic analysis revealed that the virus belonged to the newly established family *Tosoviridae* within the *Bunyavirales* order (Figure 8a). The virus, named Chatham eel tosovirus, shared 40-48.4% amino acid identity with the only other member of the family, *Turtle fraservirus-1* (accession: UKB93130.1), which was isolated from diseased turtles in Florida, USA. We attempted to uncover any other potential tosovirus sequences present in NCBI’s Transcriptome Shotgun Assembly (TSA) database and a custom database of locally downloaded and assembled fish host Sequence Read Archive (SRA) data by screening nucleoprotein, glycoprotein, and polymerase sequences from the novel eel virus and *Turtle fraservirus-1*. No such sequences were obtained.

### Effects of seawater influence on fish viromes

The possible effect of increasing seawater influence on the family-level viromes of all fish hosts studied was assessed using multivariate ordination (Figure 9). Seawater influence, increasing in salinity from freshwater, brackish to seawater, was not found to be a significant influencer of host virome composition and abundance (p = 0.17). Note that our sampling of fish host species was necessarily inconsistent across lakes due to the differing community complexity found at each sampling site (see below). When host species was also considered alongside seawater influence, it was found to be a significant virome impactor (p= 0.035). Given *Porure adintovirus-1* was found widely distributed across the smelt libraries, we repeated the multivariate analysis omitting *Adintoviridae* to assess whether this family was driving the host effect. When the analysis was repeated without this family, host species was still found to significantly influence virome composition (p = 0.042).

**Figure 9.**
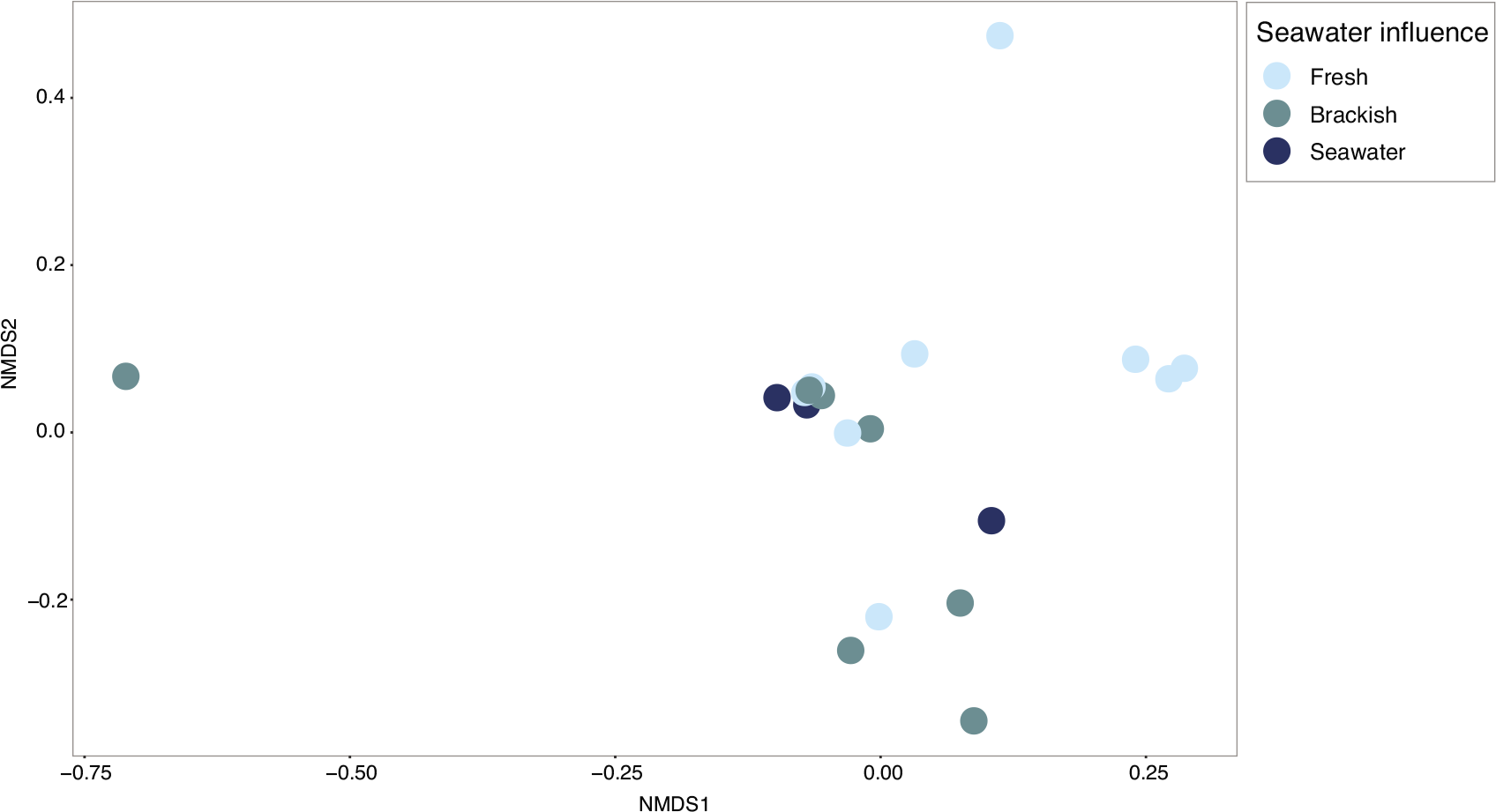
Non-metric multidimensional scaling (NMDS) plot investigating seawater influence on family-level virome composition. The 19 libraries containing fish-associated viruses were clustered by Bray-Curtis dissimilarities and coloured by the seawater influence (fresh, brackish, and seawater) of their respective sampling location. The effect of seawater influence on viromes was non-significant (PERMANOVA p = 0.171). However, host species was found to be a significant impactor of virome when it was also considered alongside seawater influence (PERMANOVA host species p = 0.035; seawater influence p = 0.266).

### Effects of ecological factors on smelt virome diversity

Due to the confounding effect of host species on virome composition when all hosts were considered (Figure 9), we further investigated the effect of seawater influence on the smelt samples containing vertebrate host-associated viruses (15/16 smelt libraries) separately from the full host set and assessed the effects of several additional ecological factors on their viromes (Figure 10). None of seawater influence (p = 0.26), community richness (p = 0.70), nor life history (p = 0.17) had significant effects on the composition of smelt viromes (Figure 10a-c). The dispersion of smelt viromes based on these three ecological factors were also assessed and was found to be non-significant in each case (ANOVA p-values = 0.57, 0.68, 0.92 respectively). Hence, smelt family-level viromes were relatively consistent regardless of their sampling environment, the number of other species present in their environment, or whether they came from diadromous or landlocked populations. Associations between life history and viral diversity were also evaluated in more detail. Viral diversity in the two types of smelt populations was measured by both richness (the total number of viral families in a sample), and Shannon diversity, which considers both the number of families (richness) and their abundances (evenness). Neither viral family richness (Figure 10d, p = 0.77) nor Shannon diversity (Figure 10e, p = 0.26) differed significantly between diadromous and landlocked smelt populations.

**Figure 10.**
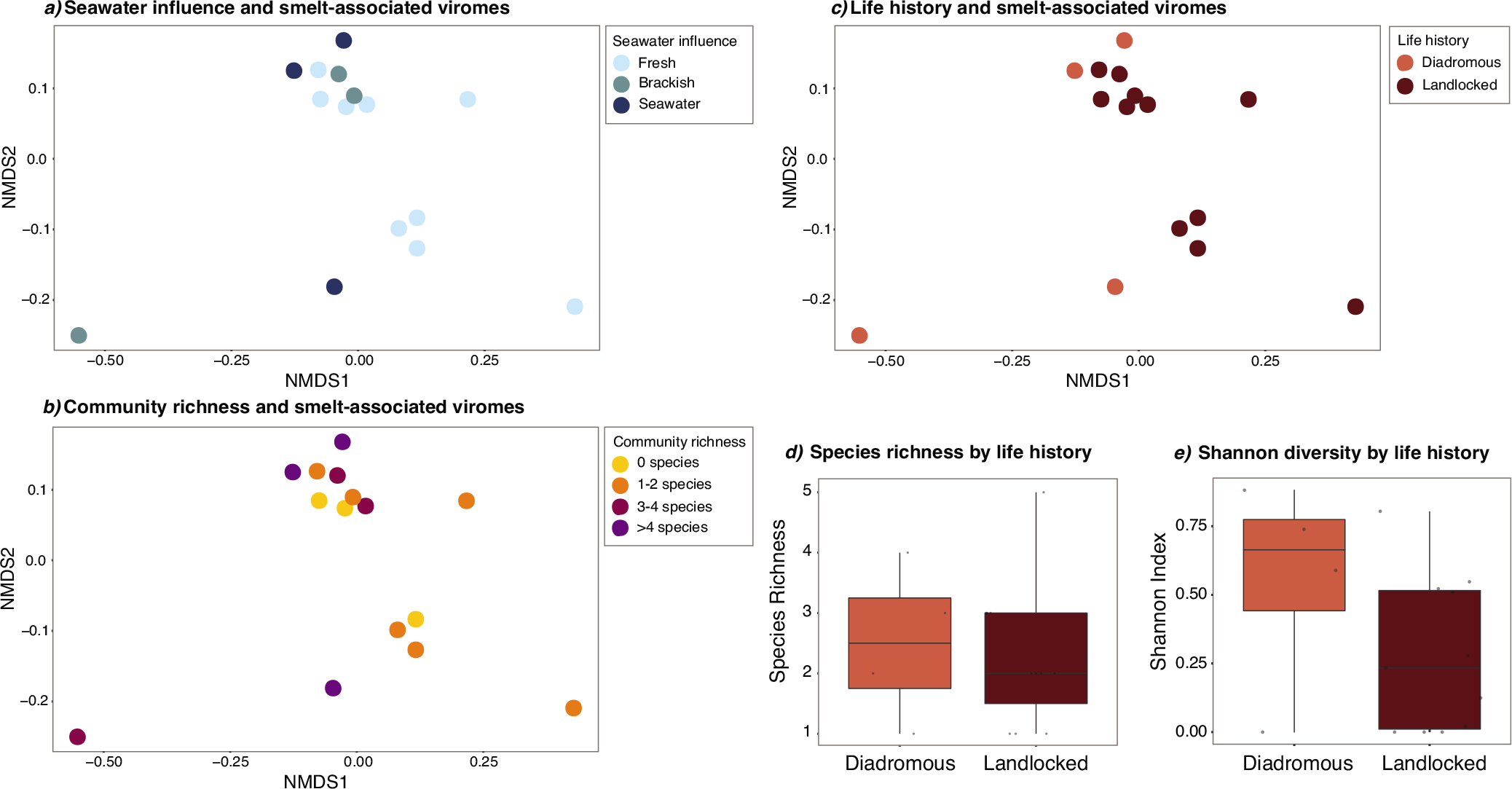
Alpha and beta diversity of smelt family-level viromes based on ecological factors. NMDS plots showing the clustering of smelt viromes based on A) seawater influence (fresh, brackish, and seawater) (PERMANOVA p = 0.259), B) community richness (either 0, 1-2, 3-4, or >4 additional species per sampling site) (PERMANOVA p = 0.697), and C) life history (diadromous or landlocked) (PERMANOVA p = 0.175) on smelt viromes. All NMDS plots were based on Bray-Curtis dissimilarities. Comparisons of alpha diversity measured by D) species richness and E) Shannon diversity were not significantly different between diadromous and landlocked (lacustrine) populations (respective Welch Two Sample t-test p-values: 0.771 and 0.258).

## Discussion

We generated metatranscriptomic data from seven freshwater fishes sampled at 16 locations across Rēkohu, Aotearoa New Zealand to investigate the diversity and distribution of their viruses. Accordingly, phylogenetic analysis identified 16 divergent novel fish viruses across 10 viral families from four of these hosts (smelt, inanga, shortfin eels, and longfin eels), and expanded the distribution of three additional viruses previously identified in smelt on Rēkohu (Perry et al., 2022). The impact of ecological factors in each sampling site on the composition and diversity of viromes of all host species (seawater influence) and smelt (seawater influence, community richness, and life histories) was also assessed.

Freshwater habitats contain roughly half of all fish species despite making up a small percentage of water on Earth (Dudgeon et al., 2006). Accordingly, we expected freshwater fish viromes to be equally diverse and likely influenced by the degree of seawater exposure in each sampling location. Studies have shown unique virome signatures and clustering of seawater and freshwater viromes (Kim et al., 2015) and that water salinity influences virome structure in aquarium systems (Kim et al., 2017). Freshwater systems were traditionally thought to harbor greater viral diversity than marine environments. However, this was primarily explored in the context of virus-bacteria associations, where a higher bacterial abundance corresponds to a higher viral abundance (Maranger & Bird, 1995), and in studies focused on disease-causing agents in fish that have typically been conducted in freshwater locations or farms due to biases induced by aquaculture practices (C. H. Kim & Leong, 1999). More recent studies on the viromes of fish from Australian rivers (Costa et al., 2021) and the Great Barrier Reef (Costa et al., 2023) have revealed diverse viromes in both freshwater and seawater systems. Our results showed that seawater influence did not impact the viromes of Rēkohu fishes, with no significant clustering of samples when host species was considered, or when smelt were evaluated separately. Such a result was unexpected given previous records of freshwater system virome diversity but could also reflect the understudied nature of fish viromes from both seawater and freshwater systems and of diadromous species in general.

Smelt were the most thoroughly sampled species in this study, allowing evaluation of two additional ecological factors – community richness and life history. Remarkably, smelt viromes were highly homogenous across all lakes, dominated by adenoviruses, adintoviruses, adomaviruses, astroviruses, and poxviruses. Previous studies have shown that fish from multispecies communities had higher viral Shannon diversity than those from single species communities (Geoghegan et al., 2021). In diverse, multispecies communities it is theorised that the dilution effect may prevent the spread of pathogens by reducing (i.e. “diluting”) the contact of infected hosts with other viable hosts from the same species (Khalil et al., 2016; Schmidt & Ostfeld, 2001). However, we observed no distinct clustering of smelt virome composition based on differing community richness between sampling sites.

Very few studies have addressed virome composition and diversity between landlocked (lacustrine) and diadromous fish populations. It might be expected that fish that complete a life cycle that involves a marine stage may be exposed to a more diverse set of viruses than those in land-locked populations. However, in a previous study of smelt and common bully (*Gobiomorphus cotidianus*) on Rēkohu and the New Zealand mainland that included three of the populations sampled in this study, it was found that lacustrine smelt populations had higher viral richness than diadromous populations (Perry et al., 2022). In contrast, with sampling of a wider set of populations we found no difference in virome composition or alpha diversity between the two life histories.

Additionally, microbial community similarity has been shown to decrease as geographical distance between communities increases (Gao et al., 2022). Using pairwise distances between lakes, we tested whether smelt virome diversity between samples followed a similar pattern, with smelt from waterbodies located further having more dissimilar viral diversity. Our analysis did not find a significant relationship between sampling location and viral diversity between samples.

While various ecological factors have shown to be significant influencers of virome composition and diversity, taxonomic relatedness of host species is consistently the strongest driving factor of fish virome composition. The phylogenetic placement of viruses identified here broadly supports the importance of virus-host co-divergence, highlighting host taxonomy and relatedness as a major factor shaping virome composition. Many of the viruses identified in these samples were related to, and expanded on, a known array of other recently discovered fish viruses (Geoghegan et al., 2021; Miller et al., 2021; Shi et al., 2018). A marine or fish origin has also been proposed for many viral families, including those found here such as the *Totiviridae*, members of the *Bunyavirales*, *Hantaviridae*, and *Flaviviridae* (Mifsud et al., 2023; Y.-Y. Zhang et al., 2022). Most importantly, viruses uncovered here fell into basal clades with other fish and ectothermic hosts. This was particularly true for DNA viruses and reflects a long history of co-divergence between viruses and their vertebrate hosts (Y. Z. Zhang et al., 2018). As a case-in-point, *Porure adintovirus-1* in smelt was widely distributed across Rēkohu with little genetic variation between the virus at different sampling sites. The wide distribution of such viruses, and the formation of distinct fish-associated clades suggests these viruses have been co-diverging closely with hosts such as *R. retropinna*, potentially before their dispersal to the island some time after it emerged <3 mya.

RNA viruses found here also fell into fish-associated clades, but many clustered with additional viruses from birds and mammals (e.g. in the *Astroviridae*), indicative of more complex evolutionary histories involving cross-species transmission events in the evolutionary past. It is being increasingly demonstrated that cross-species transmission occurs frequently at the family-level in both DNA and RNA viruses (Geoghegan et al., 2017), and particularly so in the case of RNA viruses (Holmes, 2009). Chatham eel tosovirus, for example, was identified in both shortfin and longfin eels with 99.8% sequence identity between RdRp segments at the nucleotide level. Nevertheless, these hosts are relatively closely related, having diverged less than 10 mya and it therefore may not be surprising the virus has been able to jump between these hosts (Minegishi et al., 2005). The overall long-term associations between other fish sampled here and their viruses implies that there are barriers to virus host-jumping, likely reflecting the phylogenetic distance between host species (Costa et al., 2023; Y.-Z. Zhang et al., 2018; Engelstädter & Fortuna, 2019).

The apparent lack of influence of host ecology on virome compositions and the homogeneity of smelt viromes across locations may reflect the particular composition of dsDNA viruses in these fish, coupled with the relatively young history of the island and its waterbodies. Rēkohu emerged from the Chatham Rise during the Pliocene Epoch (Stilwell & Consoli, 2012). Therefore, species such as smelt would have colonised and dispersed, forming isolated populations between the formation of the archipelago around three mya (Roberts, 1991) and the arrival of humans by 1,500 CE (Richards, 1952). Genomic analysis of smelt on Rēkohu has suggested an arrival time of around 130-570 kya followed by multiple landlocking events (Ara, unpublished data). This theorised recent dispersal of hosts may have limited the influence of fish ecology on their viromes while supporting the persistence of virus species and families in species such as smelt across multiple waterbodies. Furthermore, smelt viromes across Rēkohu share a large dsDNA virus component. Nucleotide substitution rates for dsDNA viruses have been estimated to be to the order of 10^-7^ to 10^-9^ nucleotides per site per year, akin to that of cellular genes (Domingo, 2016). Additionally, genes such as polymerases used for phylogenetic analysis have more functional constraint and often evolve more slowly than other viral genes. These factors, along with the short genomic regions analysed, may in part explain how smelt viruses sequences have remained homogenous despite their isolation and differing ecology within the proposed timeframe of colonisation and landlocking of smelt. Importantly, even when the most widely distributed smelt-associated dsDNA viruses, *Porure adintovirus-1*, was excluded from analysis, a significant host effect and lack of virome dispersion was still evident.

Interestingly, several RNA virus species, including those from the *Astroviridae* and *Totiviridae*, also appear across multiple isolated populations with little sequence divergence, despite the much higher rate of nucleotide substitution in RNA than DNA viruses (Peck & Lauring, 2018). One explanation is that these RNA virus sequences result from environmental viruses associated with the smelt, or were introduced as a result of sampling and RNA extraction and processing, and therefore denote the presence of viruses that do not directly infect fish hosts. We purposely focused our analysis on likely vertebrate-associated viruses (identified based on closest relatives and phylogenetic analyses) to help exclude common sources of viruses derived from non-fish sources (i.e., reagents, bacterial, algae, fungi, amoeba, and invertebrates) from our analyses (Asplund et al., 2019; Smuts et al., 2014). Smelt, however, are predatory, feeding on crustaceans and insects (Boubee & Ward, 1997). Members of the *Totiviridae*, the *Bunyavirales* order, *Hantaviridae*, and *Flaviviridae* are common in marine invertebrates (Y.-Y. Zhang et al., 2022), and hence it is expected that such viral families are present in our data, potentially adding uncertainty about the origin and true hosts of such virus despite being seemingly related to other fish-associated viruses. Alternatively, these RNA viruses could be spread between sites by an undetermined vector. Invertebrates, particularly those that have a flighted adult stage, may be responsible for moving certain viruses, whether vertebrate- or invertebrate-related, between locations as has been hypothesised with viruses from the *Totiviridae* (Tighe et al., 2022). Birds that prey on small fish such as smelt could also potentially spread viruses between locations (Chan et al., 2015). Ecosystem-wide virological surveys to reveal potential vectors of fish virus transmission or connectivity of dietary viruses at different tropic levels (French & Holmes, 2020) will be important in future metagenomic studies to help add confidence regarding host assignments and potential non-fish drivers of virus distribution and homogeneity between lakes.

Another notable discovery was that of a novel *tosovirus* in both shortfin and longfin eels from Lake Te Wapu (Chatham eel tosovirus). The only related virus was *Turtle fraservirus-1,* a novel virus isolated from heart cells of freshwater turtles in Florida, USA in 2007 and more recently genomically characterised in 2022 (Waltzek et al., 2022). The genome organisation of this original tosovirus resembles that of arenaviruses and hantaviruses, yet it shared no sequence similarity to any previously known virus, setting it along a deep branch within the order *Bunyavirales*. A new family, the *Tosoviridae*, was proposed to house this divergent virus. No additional tosovirus sequences were able to be recovered despite further datamining efforts. Hence, additional members of this family may be too divergent to detect using traditional sequence similarity-based approaches, coupled with inadequate sampling of their natural hosts or preferred tissue types. A future perspective may come from a more comprehensive screening of the much larger SRA database as new tools and computing power allow more efficient screening of such a database. More insight into the host range of these viruses is also warranted given their distantly related ray-finned fish and chelonian aquatic hosts, as well as its potential to cause disease. This may be of particular interest to conservation efforts as longfin eels have been assessed as being “at risk – declining” by the New Zealand Department of Conservation (Allibone et al., 2010). Currently, the tosovirus from Rēkohu represents the first virus to expand on the host range of, and solidify, this new *Tosoviridae* family and highlights the value of continuing to explore the virosphere given that even viruses with no significant sequence similarity to previously known viruses may cause severe or fatal diseases in their hosts.

While fish sampled in this study were seemingly healthy, relatives of viruses found here have been associated with disease in birds, mammals, reptiles, amphibians, and fish (Charrel & de Lamballerie, 2010; Gjessing et al., 2016; Hu et al., 2015; Schütze, 2016). For example, coronaviruses have been found as far back in the vertebrate evolutionary tree as jawless fish (Miller et al., 2021), and have been tentatively associated with disease in lamprey (Miller et al., 2021) and salmon (Mordecai et al., 2019b), with evidence that they cause disease in marine mammals, such as beluga (Schütze, 2016). We also identified a novel member of the *Poxviridae.* Poxviruses can cause diseases with mortalities of 50-90% in fish (Gjessing et al., 2016). However, without comprehensive pathology to accompany these viromics data it is unknown whether most novel viruses identified in fish cause symptomatic disease or, as more recent metatranscriptomic sampling reveals diverse viruses in the absence of apparent disease manifestation, if most are more akin to commensal microbes that are merely related to those causing notable diseases in other hosts (Harvey & Holmes, 2022).

Our study has contributed to understanding the diversity and distribution of fish viruses on Rēkohu. Although seawater influence showed some clustering of host viromes, host species was the primary factor impacting family-level virome composition. While limited cross-species transmission was observed, a highly divergent tosovirus was identified in two different, but closely related, Anguilliform species. Other viruses solidifying fish-associated viral clades, such as those DNA viruses from the *Adintoviridae* and *Poxviridae*, support the notion of long histories of host-virus co-evolution within their respective families. The homogeneity of smelt viromes irrespective of differences in life histories and environmental factors was also notable, likely driven by a diverse, largely DNA virus component. Expanding this sampling of different freshwater fauna across Rēkohu may help to reveal more definitive patterns of virus distribution and the relationships between host ecology and viromes in this unique ecosystem.

## Acknowledgements

We wish to thank Hamish Thompson for illustrating the fish host species for this project. We thank the Hokotehi Moriori Trust, Loretta Lanauze, Tom Lanauze, Cass Solomon, Maui Solomon, and Susan Thorpe for support of this research. We thank On Lee Lau and Nathan Silcock assistance during fieldwork, and landowners for permitting access to sites.

## Supplementary Material

**Supplementary Table S1.** Excel spreadsheet of full virus list.

**Supplementary Table S2.** Excel spreadsheet of all recovered segments from viruses identified in this study.

